# MYBL2 drives prostate cancer plasticity and identifies CDK2 as a therapeutic vulnerability in RB1-loss and neuroendocrine prostate cancer

**DOI:** 10.1101/2024.01.31.578216

**Authors:** Beatriz German, Jagpreet N. Singh, Marcos AdS Fonseca, Deborah L. Burkhart, Anjali Sheahan, Hannah Bergom, Katherine L. Morel, Himisha Beltran, Justin H. Hwang, Kate Lawrenson, Leigh Ellis

## Abstract

Phenotypic plasticity is a recognized mechanism driving therapeutic resistance in prostate cancer (PCa) patients. While underlying molecular causations driving phenotypic plasticity have been identified, therapeutic success is yet to be achieved. To identify putative master regulator transcription factors (MR-TF) driving phenotypic plasticity in PCa, this work utilized a multiomic approach using genetically engineered mouse models of prostate cancer combined with patient data to identify MYBL2 as a significantly enriched transcription factor in PCa exhibiting phenotypic plasticity. Genetic inhibition of *Mybl2* using independent murine PCa cell lines representing phenotypic plasticity demonstrated *Mybl2* loss significantly decreased in vivo growth as well as cell fitness and repressed gene expression signatures involved in pluripotency and stemness. Because MYBL2 is currently not druggable, a MYBL2 gene signature was employed to identify cyclin-dependent kinase-2 (CDK2) as a potential therapeutic target. CDK2 inhibition phenocopied genetic loss of *Mybl2* and significantly decreased in vivo tumor growth associated with enrichment of DNA damage. Together, this work demonstrates MYBL2 as an important MR-TF driving phenotypic plasticity in PCa. Further, high MYBL2 activity identifies PCa that would be responsive to CDK2 inhibition.

**Significance:** PCa that escapes therapy targeting the androgen receptor signaling pathways via phenotypic plasticity are currently untreatable. Our study identifies MYBL2 as a MR-TF in phenotypic plastic PCa and implicates CDK2 inhibition as novel therapeutic target for this most lethal subtype of PCa.

## Materials and Methods

### Animal ethics statement

All experiments involving mice were approved by and performed in accordance with the guidelines of the Cedars Sinai Medical Center, Institutional Animal Care and Use Committee (Animal protocol #9828).

### Cell culture, reagents, and drug response assays

Murine PBCre4:Ptenf/f:Rb1f/f knockout (DKO) 2D cell lines were generated from previously described genetically engineered mouse models (1). PBCre4:Ptenf/f:Trp53f/f knockout (PPKO) 2D cell lines were kindly provided by Dr. Zongbing You (Cunningham D., et al 2018). DKO and PPKO cell lines were cultured high-glucose DMEM and 10% FBS. For drug sensitivity assays, 500 or 350 cells/well for DKO or PPKO respectively were seeded into a 96 well plate and treated with the indicated concentrations of BGG463 (CDK2i) and CPS2 (CDK2 degrader) for 72 hours. Cell growth was assessed by CellTiter-Glo (Promega) assay and Cristal violet assay according to manufactures instructions. Cell viability/death was determined by Click-iT Plus Edu and PI staining (Thermofisher).

## Data Availability Statement

The DKO and PPKO Ctl and *Mybl2* KO spheroids were culture 21 days and RNA was extracted using Trizol protocol according to the manufacturer’s instructions. RNA-seq data were aligned with STAR to mouse reference genome mm10 (GRCm38), quantified using RSEM and normalized. Gene set enrichment analysis (GSEA) was performed in GenePattern (genepattern.org). Genotypes were compared as described, using 10,000 gene set permutations to generate normalized enrichments scores, with FDR q-value <0.25 considered significant.

Use of previously published RNA-seq data included murine prostate cancer (1,2) and were obtained from Gene Expression Omnibus (Accession no. GSE90891 and GSE92721). Human prostate cancer gene expression data from TCGA-PRAD (408 tumor samples) were obtained from the NCI Genomic Data Commons Data Portal. Stand Up to Cancer prostate dataset were obtained from the cBioPortal (3,4). LUCaP gene expression data was obtained from the Gene Expression Omnibus (Accession no. GSE126078). Human gene expression dataset for CRPC-Ad and CRPC-NE patients (5) was obtained through dbGaP (dbGaP phs000909.v.p1).

### ATAC-sequencing

ATAC-seq experiment was performed as described (6). In brief, cells were collected by incubating in trypsin for 5 min at room temperature and subsequent centrifugation at 592g for 5 min at 4 °C. Fifty-thousand cells were used for tagmentation by incubating in 50 μl of 1× THS-seq buffer (25 μl 2× THS buffer (66 mM Tris acetate, pH 7.8, 132 mM potassium acetate, 20 mM magnesium acetate and 32% dimethylformamide), 5 μl 10× Digitonin, 2 μl Illumina-TDE1) for 20 minutes at 37 °C. To stop the tagmentation reaction, an equal volume of 2× Tagmentation Stop Buffer (10 mM Tris-HCl (pH 8.0), 20 mM EDTA (pH 8.0)) was added to the reaction and incubated for 10 min on ice. For cell lysis, an equal volume of 2× lysis buffer (100 mM Tris-HCl (pH 8.0), 100 mM NaCl, 40 μg ml−1 proteinase K, 0.4% SDS) was added to the tagmentation mix and incubated at 65 °C for 15 min. The tagmented DNA library was purified in 20 μl buffer EB using Qiaquick PCR purification kit (Qiagen). Number of amplification cycles and library quantification was done as described53. Paired- end sequencing was performed on an Illumina NextSeq 500.

### ChIP-sequencing analysis

ChIP-seq experiment was performed as previously described^1^. In brief, fresh-frozen prostatic tumor tissue was pulverized (Cryoprep Pulvrizer, Covaris), resuspended in PBS + 1% formaldehyde, and incubated at room temperature for 20 minutes. Fixation was stopped by addition of 0.125 M glycine (final concentration) for 15 minutes at room temperature, then washing in ice cold PBS + EDTA-free protease inhibitor cocktail (PIC; #04693132001, Roche). Chromatin was isolated from biological triplicates by the addition of lysis buffer (0.1% SDS, 1% Triton X-100, 10 mM Tris-HCl (pH 7.4), 1 mM EDTA (pH 8.0), 0.1% NaDOC, 0.13 M NaCl, 1X PIC) + sonication buffer (0.25% sarkosyl, 1 mM DTT) to the samples, which were maintained on ice for 30 minutes. Lysates were sonicated (E210 Focused-ultrasonicator, Covaris) and the DNA was sheared to an average length of ∼200-500 bp. Genomic DNA (input) was isolated by treating sheared chromatin samples with RNase (30 minutes at 37°C), proteinase K (30 minutes at 55°C), de-crosslinking buffer (1% SDS, 100 mM NaHCO3 (final concentration), 6-16 hours at 65°C), followed by purification (#28008, Qiagen). DNA was quantified on a NanoDrop spectrophotometer, using the Quant-iT High-Sensitivity dsDNA Assay Kit (#Q33120, Thermo Fisher Scientific). On ice, H3K27ac (5 μl, catalog no. C15410196; Diagenode Diagnostics) antibody was conjugated to a mix of washed Dynalbeads protein A and G (Thermo Fisher Scientific) and incubated on a rotator (overnight at 4°C) with 1.3 μg of chromatin. ChIP’ed complexes were washed, sequentially treated with RNase (30 minutes at 37°C), proteinase K (30 minutes at 55°C), de-crosslinking buffer (1% SDS, 100 mM NaHCO3 (final concentration), 6-16 hours at 65°C), and purified (#28008, Qiagen). The concentration and size distribution of the immunoprecipitated DNA was measured using the Bioanalyzer High Sensitivity DNA kit (#5067-4626, Agilent). Dana-Farber Cancer Institute Molecular Biology Core Facilities prepared libraries from 2 ng of DNA, using the ThruPLEX DNA-seq kit (#R400427, Rubicon Genomics), according to the manufacturer’s protocol; finished libraries were quantified by the Qubit dsDNA High-Sensitivity Assay Kit (#32854, Thermo Fisher Scientific), by an Agilent TapeStation 2200 system using D1000 ScreenTape (# 5067-5582, Agilent), and by RT-qPCR using the KAPA library quantification kit (# KK4835, Kapa Biosystems), according to the manufacturers’ protocols; ChIP-seq libraries were uniquely indexed in equimolar ratios, and sequenced to a target depth of 40M reads on an Illumina NextSeq500 run, with single-end 75bp reads. BWA (version 0.6.1) was used to align the ChIP-seq datasets to build version NCB37/MM9 of the mouse genome. Alignments were performed using default parameters that preserved reads mapping uniquely to the genome without mismatches. Bam files were concatenated to sum the biological replicates of each state and bigwiggle files were calculated for comparing the different states.

### Master transcription factor analysis

Enhancer and super-enhancer (SE) calls were obtained using the Rank Ordering of Super-enhancer (ROSE2) algorithm (ref). Clique enrichment scores (CESs) for each transcription factor (TF) were calculated using clique assignments from Coltron (7). Coltron assembles transcriptional regulatory networks (cliques) based on H3K27 acetylation and TF-binding motif analysis. The clique enrichment score for a given TF is the number of cliques containing the TF divided by the total number of cliques. We incorporated ATAC-seq data to restrict the motif search to regions of open chromatin. Using the CES, we performed clustering (distance = Canberra, agglomeration method = ward.D2) considering the union set TFs. One-tailed t-test was performed to select TF specific SKO and DKO (p-value = 0.05). Motifs found by coltron were represented by sequence logos (ggseqlogo R package) using the genomic regions coordinates from Mus musculus (mm9). IGV version 2.8.13 was used to visualize normalized ChIP-seq read counts at specific genomic loci.

### *Mybl2* CRISPR-knockout cell lines

For CRISPR/Cas9-mediated knockout cell line generation in DKO and PPKO cells two step viral infection method was used. Guide RNA (gRNA) sequences CATGACCTGTCATCCGACCA and the combination of 3 guides TCTGGATGAGTTACACTACC, TTGAATCCCGACCTTGTTAA, GAGGTTTCCCAGCCGAGTCC targeting murine *Mybl2*, were cloned into the lenti-CRISPR puro (Addgene, #52963) according to the Zhang lab protocol. The scrambled gRNA sequence ACAACTTTACCGACCGCGCC and GAGCTGGACGGCGACGTAAA were used as a negative control respectively in DKO and PPKO cells. The Cas9-blasticidine vectors (Adgen96924) was used to generate the lenti-Cas9 virus. Viral infection was performed sequentially as described by the RNAi consortium (Broad Institute) laboratory protocol “Lentivirus production of shRNA or ORF-pLX clones”. After first infection with the lenti-guideRNA viruses, single clones were isolated following puromycin (2 μg/mL; Sigma-Aldrich #P8833) treatment. After second infection with the lenti-Cas9 virus, single clones were isolated following blasticidine (10mg/mL, invivo-Gen, #ant-bl) treatment.

### Growth In Low Attachment Assay (GILA)

For GILA, 500 cells from DKO Ctl and *Mybl2* KO lines, and 350 cells from PPKO Ctl and Mybl2 KO lines resuspended in 100 mL of medium were seeded in U-bottom, ultra-low attachment plates (#7007, Corning Life Sciences, Switzerland) and incubated for 5 days at 37C under 5% CO2. ATP levels were measured using RealTime-Glo^TM^ MT cell viability Assay (Promega, # PRG9713) according to the manufacturer’s instructions. The luminescent signal was detected using the SpectraMax iD3 reader with a 0.5-s measuring time.

### Cell proliferation assay

DKO and PPKO Ctl and Mybl2 KO spheroids were treated with 10 μM EdU for two hour and stained with Invitrogen Alexa Fluor 647 picolyl azide, according to the protocol for the Invitrogen Click-iT Plus EdU Alexa Fluor 647 Flow Cytometry Assay Kit (Thermofisher, #C10424), followed by staining with propidium iodide (PI) staining. Cells were then analyzed by flow cytometry using either 647 nm excitation (for Click-iT EdU Alexa Fluor 647 dye) or 493 nm excitation (for PI stain).

### In vivo therapy experiment

DKO and PPKO cells were subcutaneously implanted into syngeneic C57BL/6N mice (Jackson Laboratory). Then, 7 days (d) after the implant, mice were treated with either with 400 mg kg−1 of CPS2 (MedChemExpress, #HY-141680) or 2.5% DMSO in corn oil by intraperitoneal injection every 7 days. Tumors and mouse weight were measured three times weekly by caliper measurements. Treatment toxicities were assessed by body weight, decreased food consumption, signs of dehydration, hunching, ruffled fur appearance, inactivity, or nonresponsive behavior.

### Immunohistochemical staining’s and quantification of subcutaneously injected tumors

For all staining’s, 5 μm thick sections were cut from paraffin-embedded blocks and dried onto positively charged microscope slides. The following primary antibodies were used: Ki67 (Thermo Fisher Scientific, D3B5, 1:100), and phospho-H2AX (Cell Signaling Technology, 20E3, 1:100). Tissue sections were stained using the ImmPRESS® HRP Anti-Rabiit IgG (Peroxidase) Polymer Detection Kit (Vector Laboratories). Tissue sections were imaged using an BZ-X810 microscope (Keyence). Images were de-identified and 5 random fields per section were run through QuPath image analysis software (49) to quantify positive DAB-stained cells (as a percentage of total cells).

### Western blotting

Cell lysates for western blotting were prepared using RIPA buffer (Sigma) containing a protease and phosphatase inhibitor cocktail (Roche). 20 μg total protein was subjected to SDS-PAGE electrophoresis, then transferred to a nitrocellulose membrane. The membrane was blocked in 5% non-fat milk for one hour at room temperature, followed by primary antibody incubation (Mybl2 (MABE886, Millipore Sigma) CDK2 (2546S, Cell signaling) and CDK5 (14145S, Cell signaling)). and αGAPDH (2118S, Cell Signaling). Membranes were then incubated with anti-Mouse IgG (CST 7076S) or anti-Rabbit IgG (CST 7074S) HRP-linked antibodies diluted at 1:10,000. The immunostaining was visualized using a chemiluminescent substrate kit (ThermoFisher Scientific).

## Introduction

Second-generation androgen deprivation therapies (ADT) have provided significant life-extending therapies for recurrent, or metastatic castration resistant prostate cancer (mCRPC) patients. Most recently, a subset of these mCRPC tumors progress past ADT via phenotypic plasticity that allows the cell to adopt phenotypes that no longer rely on AR expression and signaling (CRPC-AI). Genetic aberrations such as amplification of MYCN and AURKA (8) and concurrent loss of function mutations or copy number loss events in PTEN, TP53 and RB1 tumor suppressor genes (9) are associated with CRPC-AI. These tumors exhibit multilineage cell identity that includes neuroendocrine features, a stem or basal cell-like phenotype, altered kinase signaling, altered master regulator transcription factor (MR-TF) expression, and epigenetic alterations. Currently, there are no therapeutic options to provide these patients with durable response and it is crucial that molecular mechanisms representing actionable therapy targets are identified.

The MYB proto-oncogene like 2 (MYBL2) is one of the three members of the MYB transcription factor family along with MYB and MYBL1 (10). The principal functions of MYBL2 are the regulation of cell proliferation (11), cell survival and apoptosis (12), cell cycle progression, and the control of cyclin dependent kinase (CDK) expression and activity which makes MYBL2 an essential component of the DREAM multiprotein complex (Dimerization partner, RB-like proteins, E2F2 and MuvB core) (13–15). The DREAM complex is frequently affected in cancer and the overexpression of MYBL2 has been observed in various aggressive subtypes of tumors and associated with poor clinical prognosis (16–20). MYBL2 is highly regulated at both transcriptional (E2F1,2,3) and post-transcriptional (CDKs) (21) as well as by their catalytic partners including cyclin A and E-CDK2 complex which is required during S phase. Outside of its more canonical functions, MYBL2 expression is found to be significantly enriched in stem cells or cells undergoing reprogramming when compared to somatic cells (22,23) implicating a role for MYBL2 as a pluripotency gene. More recently, it was shown that MYBL2 is a chromatin-bound partner for known pluripotency factors OCT4, NANOG, and SOX2 (24). In addition, knockout of *Mybl2* in mice failed to develop past the blastocyst stage which phenocopied mice exhibiting *Sox2* or *Oct4* knockout. MYBL2 expression level has further been shown to be critical for reprogramming via chromatin remodeling which alters the accessibility of important transcription factors driving the reprogramming process (25). These studies collectively show that MYBL2 forms an important part of a pluripotency network.

Specific to prostate cancer, gene expression analysis from mCRPC human samples and xenografts revealed high expression of MYBL2 (11,26–28). An independent study confirmed significant increase of MYBL2 in CRPC samples, and it was demonstrated that high MYBL2 could facilitate castration-resistance by promoting YAP1 activity (29). Recently, increased gene expression of MYBL2 in hormone-sensitive prostate cancer was associated with worse clinical outcomes (26). Furthermore, knockdown of MYBL2 significantly inhibited the proliferation of PCa cell lines in the absence of androgens (27). These studies do provide evidence for a role of MYBL2 driving resistance to androgen receptor signaling inhibition.

In the present study, using genetically engineered mouse models and human gene expression data, we demonstrate the enrichment of MYBL2 expression and function in prostate cancers displaying phenotypic plasticity when compared to their adenocarcinoma counterparts. Moreover, loss of *Mybl2* decreased signaling pathways related to pluripotency and proliferation highlighting the important role that MYBL2 plays in orchestrating PCa cell stemness. MYBL2 is currently not druggable, so using a MYBL2 gene signature we identified CDK2 as a potential therapeutic target for phenotypic plastic PCa. CDK2 inhibition phenocopied genetic loss of Mybl2 and significantly reduced tumor growth. Together, these data indicate that CDK2 inhibition represents a novel therapeutic approach to treat prostate cancers progressing on AR signaling inhibition and exhibiting phenotypic plasticity that includes increased expression and function of MYBL2.

## Results

### MYBL2 as a putative master-regulator transcription factor (MR-TF) in murine prostate cancer deficient for Rb1

It has been previously demonstrated in human PCa patients that chromatin remodeling can govern tumoral phenotypes (30–32). Specifically, *de novo* expression and utilization of super-enhancers (SE) identified by acetylation of histone H3 lysine 27 (H3K27ac) are demonstrated to enable activation of novel transcriptomes driving progression of PCa (33,34). To identify novel candidate MR-TFs we employed a motif-based TF connectivity approach analysis using Coltron (7) from PCa genetically engineered mouse models (GEMMs). This approach scans for TF motifs inside regions of open chromatin (defined by genotype-specific ATAC-seq profiles) nestled within SE to identify TF network “cliques” (**Figure 1A**). Factors are only nominated as candidates if their promoter is active (determined by H3K27ac), and if there is evidence of autoregulation defined by their cognate motif being present within the associated SE (**Figure 1A**). Focusing on the comparison between GEMM derived prostate tissue with specific deletion of *Pten* (SKO) and *Pten:Rb1* deletion (DKO) we identified candidate MR-TFs exclusive in DKO prostate samples (**Figure 1B, 1C and supplementary table S1**). A total of 27 TF candidates including known neuroendocrine factors *Foxa2*, *Sox2, Sox1, Insm1*, *Hes6* and *Ascl1*, showed clique enrichment in DKO, whereas *Etv6, Tcf7l2, Gata3* and *Klf9* had higher clique enrichment in SKO tissues (**Figure 1B and 1C**). Both SKO and DKO samples exhibited enrichment of H3K27ac at the SE region of *Foxa1* (**Figure 1C**) in line with previous publications which demonstrate that FOXA1 drives PCa initiation, progression and plays an important role in PCa phenotypic plasticity (35,36).

**Figure. 1.**
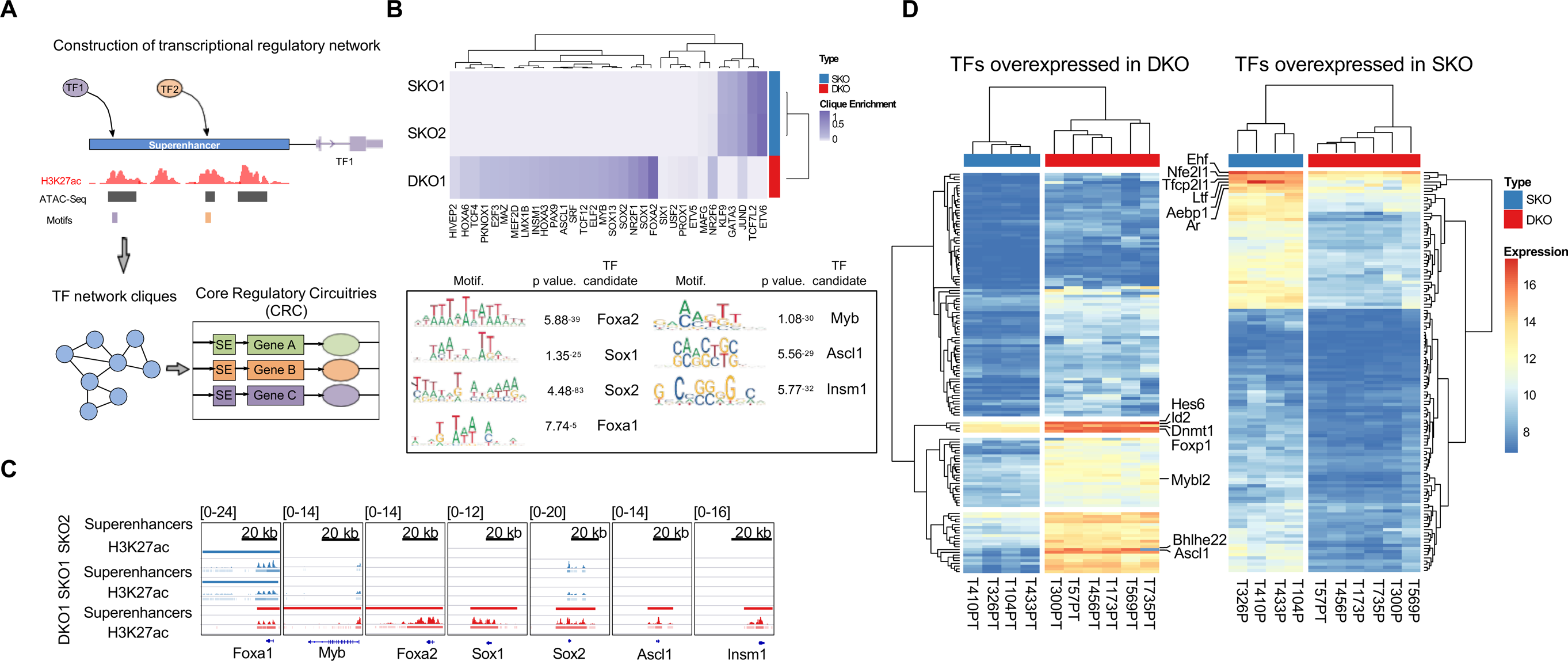
MYBL2 is a master-regulator transcription factor (MR-TF) in murine prostate cancer deficient for Rb1. **A.** Schematic representation of the Coltron algorithm procedures for construction of transcriptional regulatory networks. **B.** Candidate master transcription factors in SKO and DKO based on regulatory clique enrichment scores (range 0-1, two sided t-test p-value <0.05) and nucleotide motif logos for top candidate factors relative size of each base indicates the relative frequency at that position. Foxa2, Sox1, Sox2, Myb, Ascl1 and Insm1 motifs with the more significant P value for more abundant motifs within H3K27ac-bound chromatin were the most enriched in DKO sample. **C.** ChIP-seq tracks of H3K27ac signal at super-enhancers regions for Foxa1, Myb, Foxa2, Sox1, Sox2, Ascl1 and Insm1 in SKO (blue) and DKO (red) samples. **D.** Heat maps representing the expression profile of 128 TFs overexpressed in SKO samples (FDR </= 1-% log2 fold change >2, clustering method = “complete”, clustering distance = “Euclidean”) and 139 overexpressed DKO samples (FDR </= 1-% log2 fold change >2, clustering method = “complete”, clustering distance = “Euclidean).

The TFs overexpressed in SKO included *Ar, Ehf* and *Nfe2l1* (**Figure 1D**). Both *Ar and Ehf,* are well known as important TFs in epithelial differentiation, which downregulation promotes epithelial-mesenchymal transition (EMT) and cell dedifferentiation (37,38).

### MYBL2 expression and activity is enriched in human NEPC and NE-like mouse models

To help identify novel MR-TF’s we analyzed publicly available RNA-seq data from GEMMs (1), PDXs (39) and human prostate cancer patients (30) (**Figure 2A-C**). The integration of the differential gene expression (DEG) of the three datasets (comparing DKO versus SKO tumors, human NEPC versus adenocarcinoma, and the LuCaP NEPC verse adenocarcinoma PDX samples generated a list of 1516 common genes (**Figure 2A and supplementary table S2**). Gene set enrichment analysis (GSEA) indicated that the *fischer_dream_target* genes as the most enriched gene set in NEPC samples (**Figure 2A**). *Mybl2* is the only member of the Myb family regulated by the DREAM complex. Moreover, we found that *Mybl2* expression level and function had stronger correlation in NE-like models, including DKO, PPKO (*Pten:Trp53* KO) and DKO-CR (GEMMs with the addition of a spontaneous *Trp53* mutation) than the SKO (*Pten* KO) model (**Figure 2B**). Looking over the spectrum of disease progression in human PCa samples, NEPC patient tumors represented the most enriched sample set for increased *MYBL2* expression. Of interest, a subset of patient CRPC (adenocarcinoma) samples did exhibit medium to high levels of *MYBL2* (**Figure 2C and 2D**). While increased MYBL2 expression in PCa are linked to resistance to ARSI, it remains unknown if MYBL2 mechanistically increases the potential for PCa to activate alternate lineages within the phenotypic plasticity spectrum when progressing on AR targeted treatments. Further, whether MYBL2 stands as a biomarker of disease progression and can indicate therapy options are questions currently not answered.

**Figure 2.**
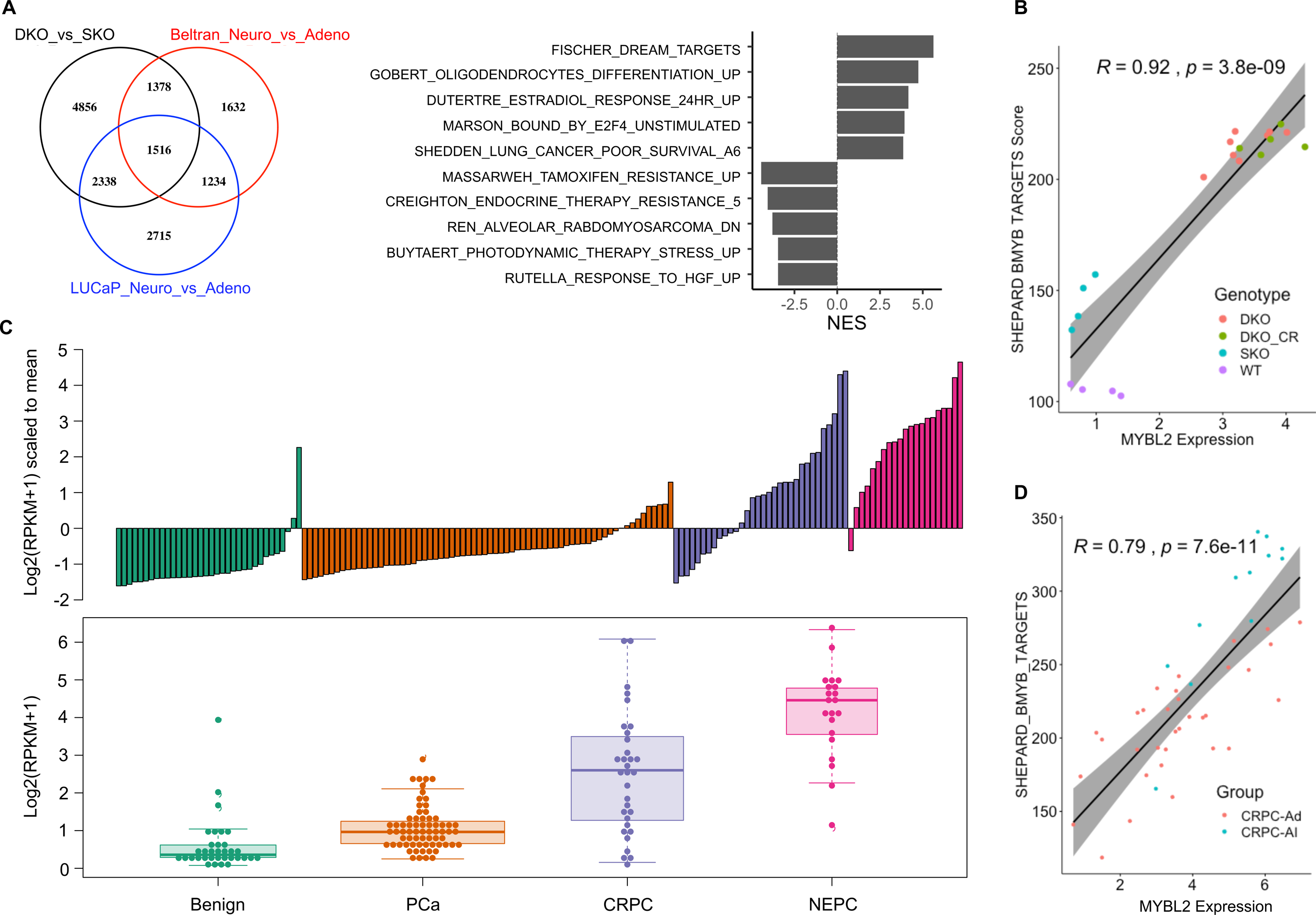
MYBL2 expression and activity is enriched in human NEPC and NE-like mouse models. **A.** Venn diagram integrating the differential gene expression (DEG) list from PBCre4:Ptenf/f (SKO) versus PBCre4:Ptenf/f:Rb1f/f (DKO) mouse models, human NEPC versus adenocarcinoma samples and LuCaP patient-derived xenografts (PDXs) obtained from human NEPC versus adenocarcinoma patients show that the 3 independent data sets shared 1516 genes (left). Gene set enrichment analysis (GSEA) of the 1516 shared genes indicates that the fischer_dream_targets genes may be involved in the regulation of CRPC-AI development (right). All the pathways listed are statistically significant with p<0.05. **B**. RNA-Seq analysis from different GEMMs (SKO, DKO, DKO_CR and WT) indicates that a MYBL2 signature and expression has stronger correlation in DKO and DKO_CR models. R=0.92, p=3.8e-09. WT (wild type), DKO-CR (PBCre4:Ptenf/f:Rb1f/f:Trp53^ι1^). **C.** Bar and box plot summarizing the MYBL2 expression level in human PCa patients at different stages of the disease progression. Adjusted P value <0.05. **D.** MYBL2 function is upregulated in CRPC-AI. R=0.79, p=7.6e-011. CRPC-Ad (castration resistant prostate cancer, adenocarcinoma), CRPC-AI (castration resistant prostate cancer, androgen indifferent).

### Mybl2 supports stemness and cell fitness in prostate cancer models

Given the role of MYBL2 in proliferation (40), genome stability (41–43) and reprograming (24), we sought to understand cellular consequences when *Mybl2* expression was lost in murine PCa cell lines. First, we used the DepMap database (44) through the Broad Institute to determine the dependency of the MYB genes in human PCa cell lines. In line with our patient data, the human neuroendocrine cell line H660 exhibited the highest expression of all MYB genes. Interestingly, MYBL2 was the only family member that represented a genetic dependency in all cell lines (Fig. 4A). We validated genetic dependency by using independent murine PCa cell lines generated from GEMMs representing phenotypic plasticity. We observed that CRISPR/Cas9 deletion of *Mybl2* (**Figure 3B, Supplementary Figure 1A**) didn’t disrupt the overall growth potential of these cells in anchorage dependent 2-dimensional (2D) culture conditions (**Supplementary Figure 1B**). However, in anchorage independent 3-dimensional (3D) culture conditions through use of a GILA assay, we observed a significant loss in spheroid formation ability. In addition, loss of Mybl2 also repressed cell growth via significant loss in DNA replication (**Figure 3C-E**).

**Figure 3.**
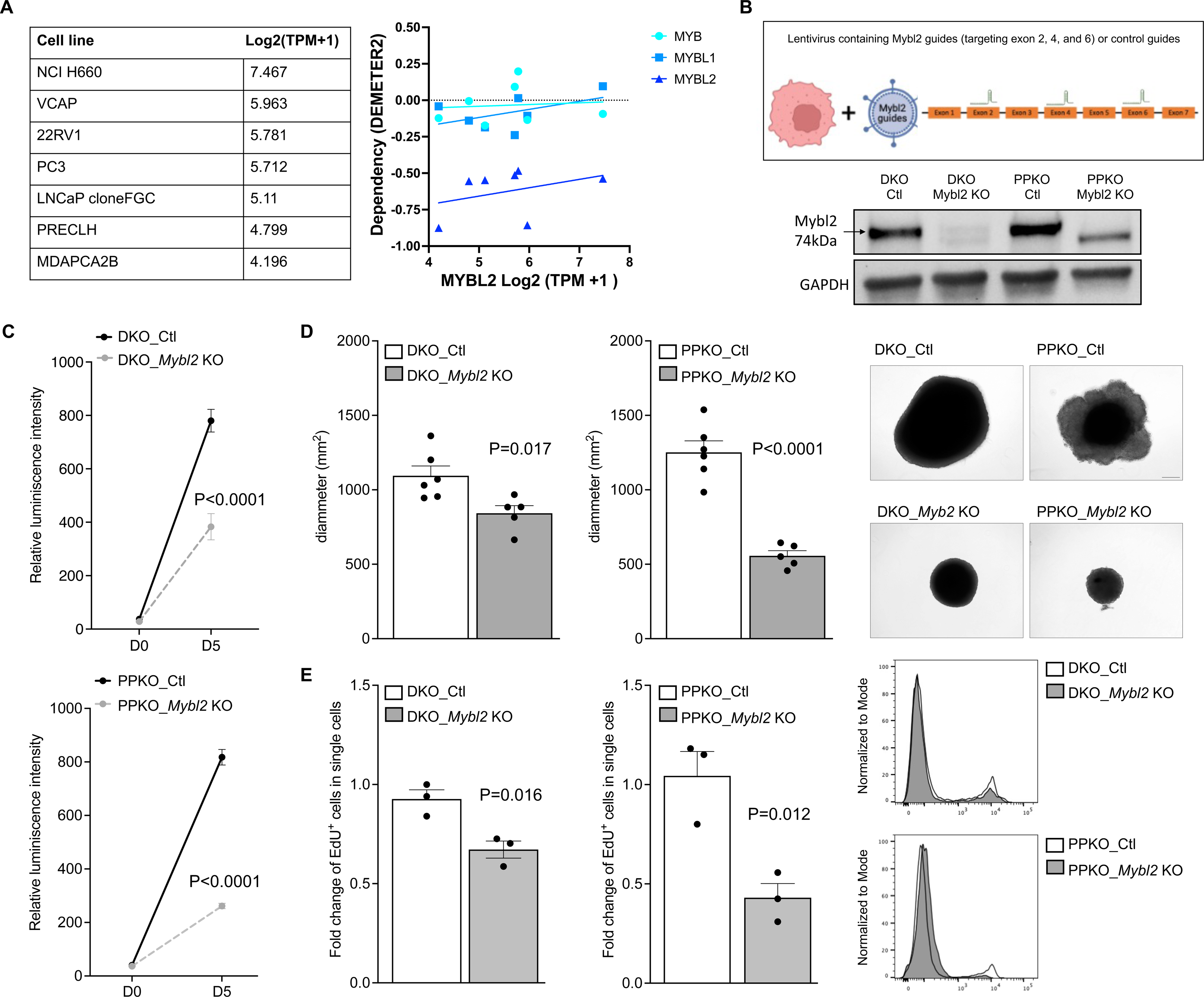
Mybl2 supports tumoral proliferation and promotes self-renewal in prostate cancer cell line models. **A.** Normalized gene expression of *MYBL2* in human prostate cancer cell lines. Demeter gene dependency of MYB family members across human prostate cancer cell lines indicating *MYBL2* as a significant dependency compared to other family members. **B.** Schematic representation of the DKO Ctl and DKO *Mybl2* KO and PPKO Ctl and PPKO *Mybl2* KO cell line generation using two step CRISPR/Cas9 method. KO: knockout. Western blot indicating Mybl2 protein level in the DKO Ctl, DKO *Mybl2* KO, PPKO Ctl and PPKO *Mybl2* KO generated cells determined by wester blotting. Ctl: Control. **C.** Growth kinetics representation using relative luminescence intensity of DKO Clt versus DKO *Mybl2* KO spheroids and PPKO Clt versus PPKO *Mybl2* KO spheroids cultured from D0 (day 0) to D5 (day 5) in a low attachment round bottom 96 well plate. **D.** Spheroid diameter and pictures of the DKO versus DKO *Mybl2* KO and PPKO versus PPKO *Mybl2* KO cells 21 days post-seeding of 500 or 350 cells/well respectively. **E.** Flow cytometry analysis of the EdU positive cells in DKO versus DKO *Mybl2* KO and PPKO versus PPKO *Mybl2* KO cells after 2hr of EdU-labeling in vitro. Data represent the fold change and histograms of EdU^+^ cells.

**Figure. 4.**
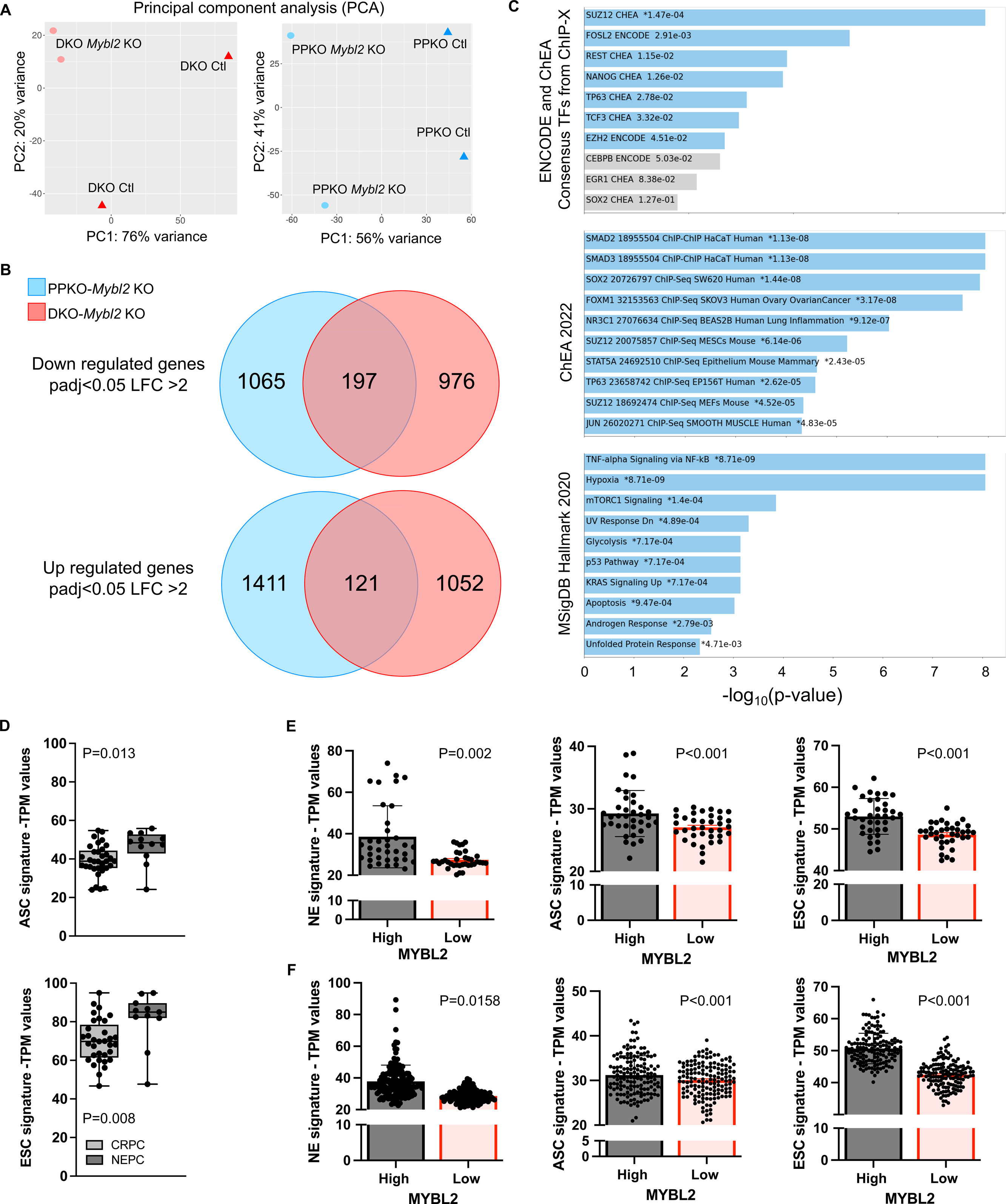
Increased Mybl2 expression supports stem related gene signatures in murine and human prostate cancer. **A**. Two-dimensional principal component analysis (PCA) visualization of bulk RNA-seq analysis performed in DKO Ctl versus DKO *Mybl2* KO cells (left) and PPKO Ctl versus PPKO *Mybl2* KO cells. **B.** Venn diagram representation of down and up regulated genes from the DEG analysis between DKO *Mybl2* KO versus DKO Ctl cells and PPKO *Mybl2* KO versus PPKO Ctl cells showed that both cell lines shared 197 down and 127 up regulated genes. The P adjusted value used to calculate the DEG was <0.05 and the LFC was < or > 2. **C.** Bar chart visualization of the top ten enriched terms and their p-values using ENCODE and ChIP-x Enrichment analysis (ChEA), ChEA 2022, and Hallmarks 2020 databases using the 197 down regulated genes shared between DKO *Mybl2* KO and PPKO *Mybl2* KO cells. All pathways listed were statistically significant with p<0.05. **D.** Normalized gene expression values indicating that human NEPC acquires increased adult stem cell (ASC) and embryonic stem cell (ESC) gene signatures compared to prostate adenocarcinoma. **E.** Normalized gene expression values for TCGA PRAD and **F.** SU2C CRPC patient samples show high *MYBL2* expression (upper 25% of samples) is associated with significantly increased expression of gene signatures for NEPC (NE), ASC, and ESC signatures when compared to patient samples with lowest *MYBL2* expression (lower 25% of samples).

Gene expression analysis by RNAseq revealed the loss of *Mybl2* expression induced a dramatic change in overall transcriptome profiles **(Figure 4A**). Furthermore, comparing the DEG between *Mybl2* KO with their respective controls we found 197 shared downregulated and 121 up regulated genes between both cell lines (**Figure 4B, Supplementary table S3**). In line with our observations in 3D culture, gene enrichment analysis demonstrated a significant repression of gene sets associated with pluripotency, stemness, and reprogramming including Fosl2, Ezh2, Suz12, Rest, Sox2, Foxm1, Tp63 and Nanog (45–47) (**Figure 4C**). We expanded our analysis to first ask if human NEPC exhibited increase in stem cell gene expression signatures. As expected, this held true with NEPC patient samples demonstrating a significant increase of both embryonic and adult stem signatures when compared to CRPC-adenocarcinoma patient samples (**Figure 4D**). Based on the initial observation from figure 2C where a subset of CRPC-adenocarcinoma samples displayed increased gene expression for *MYBL2*, we extended this analysis to adenocarcinoma patient samples from the TCGA (**Figure 4E**) and SU2C (**Figure 4F**) datasets. Comparing the *MYBL2* expressing top and bottom 25% of samples we observed that high *MYBL2* expression was associated with increased neuroendocrine, and adult and embryonic stem cell gene signatures. These data indicate that Mybl2 promotes pluripotency and stemness networks associated with phenotypic plasticity and can stand as a genetic dependency in prostate cancer models.

### Mybl2 transcriptional activity identifies CDK2 as a novel therapy target for phenotypic plastic prostate cancer

To evaluate the contribution of *Mybl2* expression in tumoral growth *in vivo*, mice were injected subcutaneously with DKO or PPKO cells with *Mybl2* wildtype (Ctl) or *Mybl2* KO cells. Loss of Mybl2 expression significantly reduced the potential for overall *in vivo* tumor formation and growth (**Figure 5A and 5B**). These data highlight MYBL2 as a promising therapeutic target, but currently no direct targeted therapies towards MYBL2 are known. Therefore, we used an established MYBL2 gene signature (48) to identify and nominate candidate drug targets as a novel therapeutic direction using the cancer therapeutics response portal for PCa patients with overexpression of MYBL2 and/or increased MYBL2 activity (**Figure 5C**). From this analysis, numerous candidate druggable pathways were nominated including PI3K/mTOR signaling, RTK signaling, protein stability and degradation, and DNA replication. Of most interest to us because of potential novelty for treating phenotypic plastic PCa was the cell cycle related cyclin-dependents kinases (CDK) which included CDK2, CDK5, and CDK7. Based on information gained from this analysis and the success of CDK inhibitors in other cancers (49,50), we applied the DEMETER computational method (44) to infer the dependency of these three CDK genes in the human PCa cell lines mentioned in figure 3. We found that CDK2 demonstrated consistent dependency within all PCa cell lines including the H660 cell line (**Figure 5C**). In line with these dependency data, we observed that NE-like GEMMs and NEPC patients had significantly higher CDK2 expression when compared to their adenocarcinoma counterparts (**Figure 5D**).

**Figure 5.**
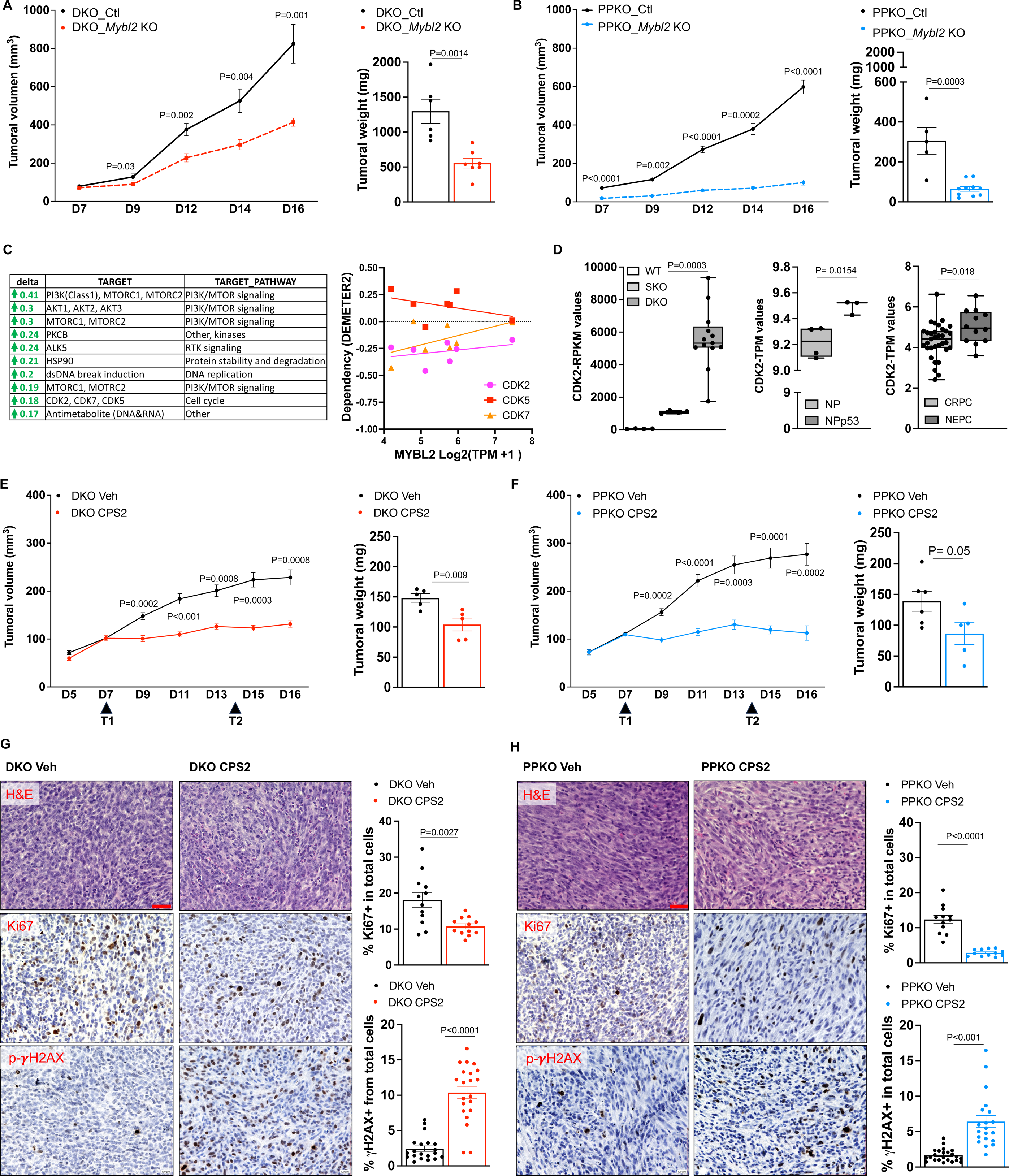
Mybl2 transcriptional signature identified CDK2 as a novel therapy target for NEPC. **A**. Tumoral growth curve of murine DKO Ctl and DKO *Mybl2* KO cells and **B.** PPKO Ctl cells and PPKO Mybl2 KO cells in NOD SCID mice. Tumor growth was monitored by serial caliper measurements every second day for 16 days (D16). Tumor weight was measured at D16 after dissection (n=5 mice per treatment group (+/-1SEM). **C.** Top responses from the cancer therapeutics response portal analysis using a MYBL2 gene signature. Demeter gene dependency of CDK family members 2, 5, and 7 across human prostate cancer cell lines indicating as a significant as outlined in Fig. 3A. **D.** Box plot graphs summarizing the RNA-Seq analysis from published prostate cancer GEMMs (first and second graphs, including WT, SKO, DKO, NP:Nkx3.1^CreERT2^:Ptenf/f and NPp53:Nkx3.1^CreERT2^:Ptenf/f:Trp53f/f) and human samples (third graph, including CRPC and NEPC samples) indicate CDK2 upregulation in NEPC mouse models and human patients. **E**. Tumoral growth curve of murine DKO cells in C57BL/6N and **F.** PPKO cells in C57BL/6N mice and treated with either vehicle control or the CDK2 PROTAC - CPS2. Mice were treated every 7 days (D7 and D14) with CPS2 (400mg/kg) by intraperitoneal (IP) injection. Tumor growth was monitored every second day for 16 days (D16) by serial caliper measurements. Tumoral weight was measured at D16 after dissection (n=5 mice per treatment group (+/-1SEM). **G-H.** Example pictures of H&E, Ki-67 and p-ψH2AX IHC staining in murine DKO and PPKO tumors from the in vivo study, and corresponding quantification of the percentage of positive nuclei from total cells (n = 5 mice per treatment group, +/-1SEM).

To investigate the potential targeting of CDK2 in phenotypic plastic tumors we initially tested a CDK2 inhibitor (BGG463) (51,52) in vitro (**Supplementary Figure 2A**). Response to BGG463 was more significant in DKO and PKO murine cell lines when compared SKO cell response. While BGG463 demonstrated superior efficacy in phenotypic plastic PCa cell lines, it is shown that BGG463 can exhibit indirect inhibition of alternate kinase function not related to CDK2, particularly inhibition of T315I BCL-ABL autophosphorylation in murine models of chronic myeloid leukemia (CML) (52). For this reason and wanting to test a more efficient on-target inhibitor of CDK2, we utilized the CDK2 degrader, CPS2 (53). Because of initial superiority of response, we only tested CPS2 ability to directly degrade CDK2 in vitro using DKO and PPKO cells. Previous analysis (53) showed that CPS2 did specifically degrade CDK2, and we repeated these results in DKO and PPKO cells showing direct down-regulation of CDK2 expression, when compared to CDK5 which remained stably expressed (**Supplementary Figure 2B-E**). *In vivo* treatment of C57BL/6N WT mice harboring DKO or PPKO tumors confirmed the ability of CPS2 treatment to significantly reduce the overall growth of DKO and PPKO tumors (**Figure 5E-D**) which phenocopied the loss of Mybl2 expression (**Figure 5A-B**). Moreover, CPS2 treatment response in both DKO and PPKO tumors was confirmed by using immunohistochemistry to show that loss of tumor growth was associated with loss of overall proliferation as indicated by lower Ki67 nuclear staining, and increased DNA damage indicated by elevated nuclear staining for ψH2AX (**Figure 5G-H**).

## Discussion

Despite the advancement of available therapy options for PCa patients with mCRPC, progression is still inevitable. While most tumors that progress and maintain AR dependency, approximately 15-20% patients will progress where their tumor is indifferent to AR signaling (CRPC-AI) and activate developmental programs related to stemness and pluripotency (phenotypic plasticity). For these patients’ treatment options remain extremely limited, and it is imperative that novel therapeutic approaches are identified.

In this study, we identified MYBL2 as a putative MR-TF driving progression to CRPC-AI. These results were in agreement with previous publications where MYBL2 was characterized as a putative oncogene (54) promoting the malignant progression of tumors by controlling cancer cell proliferation, therapy resistance, and metastasis (28). While MYBL2 is implicated in CRPC-adenocarcinoma growth (55,56), no studies have examined the specific functional and clinical implications of increased MYBL2 expression in driving phenotypic plasticity in PCa. MYBL2 expression and function was increased as localized PCa progresses to CRPC-adenocarcinoma (26,57). However, our work demonstrates that MYBL2 is more significantly increased in mouse models representing phenotypic plasticity and patients with NEPC.

Given the role of Mybl2 in proliferation (40), we asked whether the loss of Mybl2 impaired the growth of NE-like murine PCa cells. Surprisingly, loss of Mybl2 in DKO cells did not inhibit overall growth of 2D cell cultures. However, using a 3D sphere-formation assay (SFA), loss of Mybl2 did significantly perturb sphere-forming efficiency involving loss of DNA replication and indicating a dependence for Mybl2 expression in maintaining a higher self-renewal capacity. Gene expression analysis further supported this by indicating that Mybl2 mediated transcriptional programs related to self-renewal and pluripotency, as well as increased stem cell signatures in NEPC patient samples and adenocarcinoma patients with highest MYBL2 expression. Our data is in line with others, where overexpression of MYBL2 is shown to inhibit neural and glial differentiation induced by retinoic acid (58). Additionally, MYBL2 is known to be critical for the development of hematopoietic stem cells (59), and coordinates self-renewal and pluripotency by regulating expression of pluripotency genes including *POU5F1*, *SOX2*, and *NANOG* in embryonic stem cells (22,60,61).

While the MYB family of transcription factors, including MYBL2, are emerging therapeutic targets in hematological and solid malignancies, no direct MYB targeting therapy has been identified. Using a MYBL2 gene signature (48) we identified CDK2 as a candidate therapeutic target. MYBL2 and CDK2 are molecularly linked as CDK2 is a transcriptional target of MYBL2 (62), and CDK2 phosphorylates MYBL2 to enhance transactivation (63). Indeed, we demonstrate that CDK2 inhibition presents a more robust anti-tumor response *in vivo* using our phenotypic PCa tumor models. Both CDK2 (64) and MYBL2 (65,66) are implicated in DNA damage repair response, however, more insight is needed to determine if there are co-dependencies requiring cooperation between CDK2 and MYBL2 for this purpose.

Together, our study provides evidence that MYBL2 is an important MR-TF driving stemness in NEPC, and this increase in transcriptional function of MYBL2 highlights that NEPC would be susceptible to inhibition of CDK2. Previously, MYBL2 mRNA expression has been suggested as a biomarker to identify myelodysplastic syndrome patients that would respond to genotoxic treatments (67). These data raise rationale that CDK2 inhibition should be candidate target taken to clinical trial to test towards the treatment of NEPC patients.

## Acknowledgements

This study was supported by funding from the Department of Defense Translation Science Award (L.E.: W81XWH-20-1-0056); Department of Defense Early Career Investigator Award (B.G.; W81XWH-22-PRCP-EIRA), and the National Cancer Institute (L.E.; R01CA207757, R01CA252468, R21CA257484). We would like to thank the Applied Genomics, Computation and Translational Core and the Biobank and Research Pathology Resource Core at Cedars-Sinai Medical Center for services towards RNA sequencing and tissue preparation.

The contents of this publication are the sole responsibility of the author(s) and do not necessarily reflect the views, opinions or policies of Uniformed Services University of the Health Sciences (USUHS), The Henry M. Jackson Foundation for the Advancement of Military Medicine, Inc, the Department of Defense (DoD) or the Departments of the Army, Navy, or Air Force. Mention of trade names, commercial products, or organizations does not imply endorsement by the U.S. Government.

## Author Contributions

Conceptualization: LE, BG Methodology: LE, BG, KL Investigation: LE, BG, KLM, DLB, HB, JNS, MASF, HB, JHH, CJS, AS, HB, KL Writing/Editing: LE, BG

## Supplement Figure Legends

**Supplement Figure 1.**
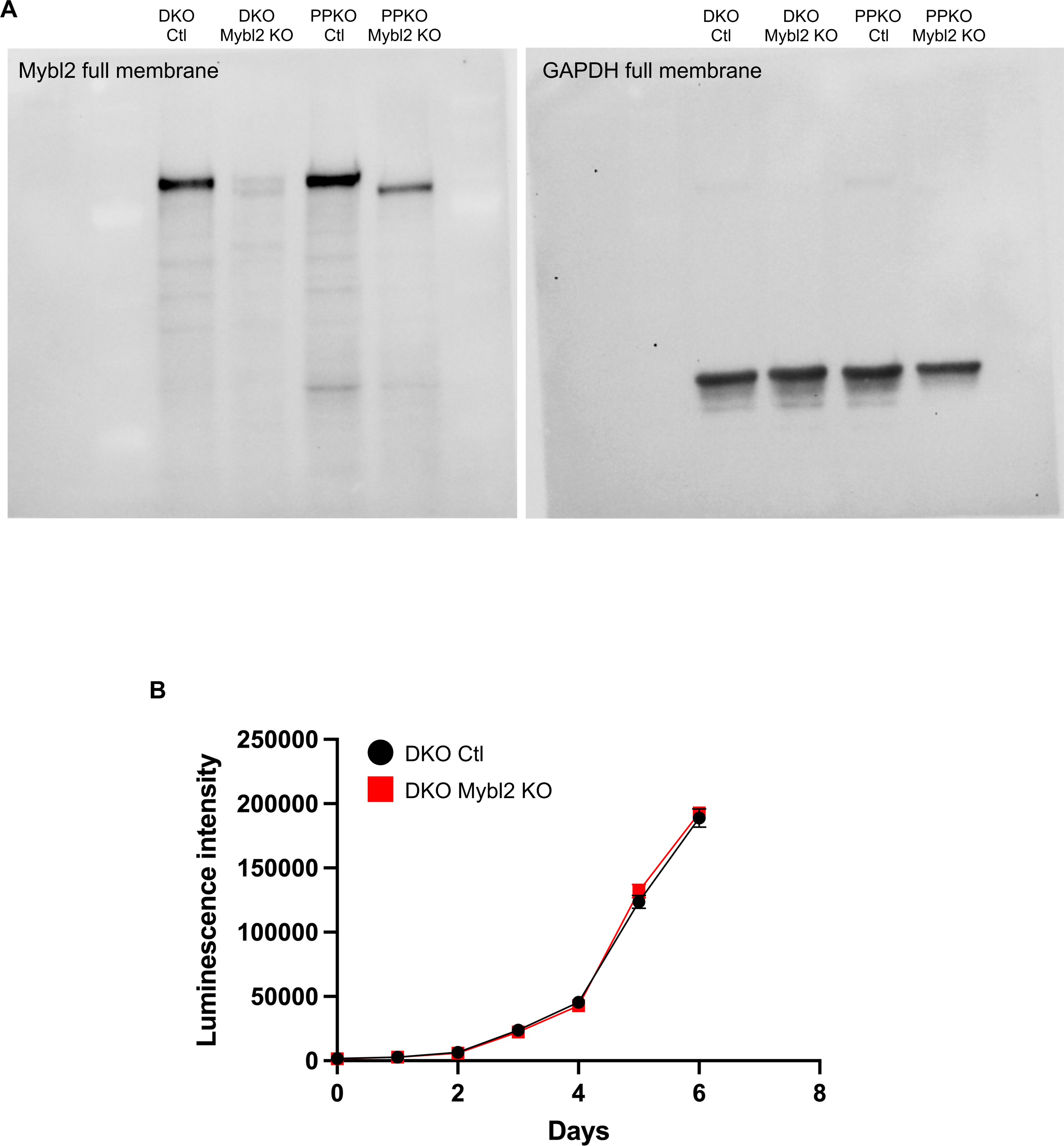
**A.** Full membrane following CRISPR targeting of *Mybl2* indicating specific targeting of our target gene. **B.** DKO-Ctl and DKO-*Mybl2* KO cells were cultured as 2-dimensional monolayers and overall cell growth was determined by CellTiter-Glo luminescent cell viability assay.

**Supplement Figure 2.**
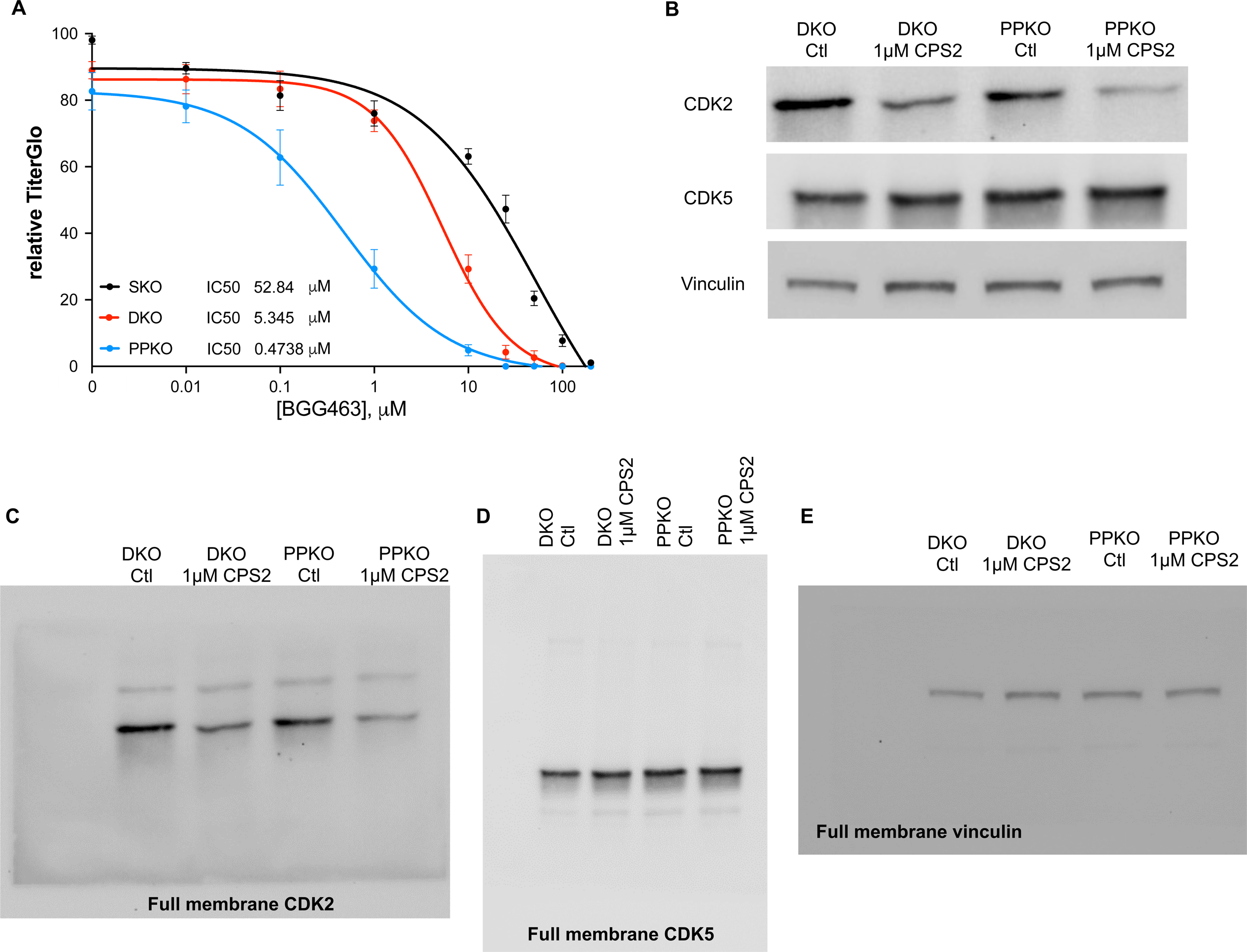
**A.** SKO, DKO, and PPKO were cultured as 3-dimensional spheroids and treated with the indicated concentrations of the CDK2 inhibitor – BGG463 over 72 hours. Cell growth was determined by CellTiter-Glo luminescent cell viability assay. **B.** Western blot analysis for Cdk2, Cdk5, and Vinculin following DKO and PPKO were treated with the indicated concentration of the CDK2 PROTAC – CPS2.

